# Optical Genome Mapping for detecting Homologous Recombination Deficiency (HRD) in human breast and ovarian cancers

**DOI:** 10.1101/2022.12.23.521790

**Authors:** Sandra Vanhuele, Youlia Kirova, Anne-Sophie Hamy-Petit, Audrey Rapinat, David Gentien, Céline Callens, Marie-Charlotte Villy, Fabien Reyal, Anne Vincent-Salomon, Alexandre Eeckhoutte, Manuel Rodrigues, Marc-Henri Stern, Tatiana Popova

## Abstract

Homologous recombination deficiency (HRD) leads to genomic instability that marks HRD tumor genome with a specific genomic scar. Present in many cancers, HRD is important to be detected as it is associated with a hyper-sensitivity to some classes of drugs, in particular the PARP inhibitors. Here, we investigated the use of structural variants (SVs) detected by the Optical Genome Mapping (OGM) technology as biomarkers to identify HRD tumors. We analyzed SVs data obtained by OGM from 37 samples of triple-negative breast cancer or high grade ovarian cancer with the known HRD status. We found that HRD cases were enriched with duplications and reciprocal translocations, while nonHRD cases were enriched with inversions. The number of translocations, defined as inter-chromosomal or intra-chromosomal rearrangements of more 5Mb were similar in HRD and nonHRD cases. We defined isolated translocations as the subset of translocations having no other translocation within 2 megabase zone around both junctions, and demonstrated that the number of isolated translocations perfectly discriminated HRD and nonHRD cases in the training series. Validation series consisting from 26 cases showed 20% false positive and zero false negative error rate, which proved isolated translocations to be 100% sensitive and 80% specific SV marker of HRD.

Our results demonstrate that the OGM technology is an affordable way of getting an insight of the structural variants present in solid tumors, even with low tumoral cellularity. It represents a promising technology for HRD diagnosis, where a single marker already gives 80% correct recognition.

## Introduction

Our cells are constantly exposed to endogenous and exogenous damage that alters DNA. Double Strand Breaks (DSB) of the DNA are the most toxic type of damage as they will lead to aneuploidy if left unrepaired before cell division. This repair can be performed using several different pathways (Ceccaldi et al. 2016; Scully et al. 2019), but the Homologous Recombination (HR) pathway is the only one able to repair *ad integrum* the genome, since it utilizes an intact copy of the broken genome region as a template, *i.e.* the sister chromatid after replication or the homologous chromosome during meiosis. Key actors in the HR pathway include BRCA1, BRCA2 and PALB2. These large proteins bind to the ends of the broken DNA to deposit on the single stranded DNA a microfilament made up of RAD51 and RAD51 paralogues. This microfilament will recognize and invade the homologous sequence to copy it. The continuity of the broken DNA will be eventually restored after resolution of the chromatid or chromosome bridges (Holliday junctions). Three other main mechanisms of repair of DSBs are the Non-Homologous End Joining (NHEJ), a process that is tolerant to nucleotide changes and insertion or deletion of bases (indel); Alternative-End Joining (Alt-EJ) / Microhomology-mediated end joining (MMEJ), a process that induces systematic small deletions with microhomologies: and Single Strand Annealing (SSA), which induces large deletions.

Homologous recombination deficiency (HRD) is an important feature to be recognized in cancer (Groelly et al. 2022). Many possible mechanisms may explain such deficiency. The main causes are mutations of key HR actors, most frequently *BRCA1* or *BRCA2*, but *PALB2, RAD51* paralogs (*RAD51C, RAD51B* and possibly *RAD51D*) mutations also play a significant role. These mutations could be inherited in hereditary breast and ovarian cancer syndrome (HBOC) or somatic. The second wild-type allele of the respective HR gene is inactivated in HRD tumors by deletion of the region, or more rarely by somatic mutations. Epigenetic inactivation of HR genes, mainly *BRCA1* and *RAD51C,* by hypermethylation of the promoter region of these genes is also a frequent mechanism of HRD. HRD is found in approximately half of triple negative (hormone receptors negative, no over-expression of HER2) breast carcinomas (TNBC) and of high-grade serous ovarian carcinomas, and in many other solid tumors at lower frequencies (Riaz et al. 2017; Knijnenburg et al. 2018).

Discovered in cellular models (Bryant et al. 2005; Farmer et al. 2005), the hypersensitivity of HRD tumors to PARP inhibitors was demonstrated in clinical trials (Mirza et al. 2016; Ray-Coquard et al. 2019). Thus, establishing the HR status of tumors, especially for high-grade serous ovarian carcinomas, is highly clinically relevant. Different methods have already been developed to diagnose HRD. Since this deficiency is due to the inactivation of HR genes target-sequencing of these genes is the most direct approach. Despite its large use in clinics, some limitations were identified, such as interpretation of variants of unknown significance and of variants in genes with more distant role in HR (for example *ATM*, *CDK12…*), the lack of detection of *BRCA1* and *RAD51C* methylation, which are major players of HRD, and of course the inability to detect HRD of unknown origin. The current strategy consists in measuring patterns of somatic alterations of the cancer genome (mutations or structural rearrangements) directly due to the DNA repair defect. The first generation of HRD signatures, extracted from SNP-arrays, includes large genomic changes (LST; Large Scale Transition), the amount of Loss of Heterozygosity (LOH) as well as the Telomeric Allelic Imbalance (TAI) (Abkevich et al. 2012; Birkbak et al. 2012; Popova et al. 2012). Those signatures are presently the FDA-approved tests used in clinics. Large sequencing panels or Whole Genome Sequencing (WGS) allow the measure of the Single Base Substitution (SBS) signature 3 shown to be associated with *BRCA1/2* deficient tumors (Polak et al. 2017). WGS allows not only the recognition of SBS3, but also the exhaustive description of rearrangement signatures, some of them being strongly associated with HRD, such as RefSig R5 characterized by deletions smaller than 100kb. RefSig R3, characterized by a number of tandem duplications (TD) smaller than 100kb has also been shown to be associated specifically with *BRCA1*-mutated tumors (Nik-Zainal et al. 2016; Davies et al. 2017). This WGS approach and the associated diagnostic pipeline HRDetect are highly performant but remain costly in terms of sequencing and data storage, and demanding in terms of bioinformatics.

We here explore Optical Genome Mapping (OGM) as an alternative from WGS for its ability to detect HRD. OGM is an affordable genome-wide visualization technique able to detect structural rearrangements. It is non sequencing based and does not employ mechanical forces to destroy the DNA. Molecules larger than 150kb are directly extracted and labeled with fluorophore tags approximately 15 times every 100 kb. Labelled DNA is then linearized on a chip where nano channels are engraved in order to image unique large DNA molecules on the Saphyr genome imager. Molecules are then aligned together to create a consensus optical map that is automatically compared to a reference map in a genome wide fashion and for which any deviation in the labelling pattern or molecule coverage indicates the presence of a structural variant (SV) or a copy number variant (CNV). The entire genome has a high coverage, usually beyond 300X, allowing to detect even rare variants. Evaluation of patterns compared to a reference is then used to detect structural variants. OGM was already described detecting SVs in hematopoietic and solid tumors (Goldrich et al. 2021; Neveling et al. 2021; Shim et al. 2022). One application included the attempt to reproduce HRD score using OGM (Sahajpal et al. 2023). Here we are testing possibility of building HRD classifier based on direct output from OGM in TNBC and HGOC.

## Materials and Methods

### Whole Genome Sequencing

Whole Genome Sequencing (WGS) was performed with Illumina NovaSeq 6000 paired-end technology and selected read length was 2 x 150bp. Resulting fastq files were aligned to the reference genome hg38 using Burrows-Wheeler Aligner v0.7.15. Duplicates were marked with Sambamba and removed with Samtools. Resulting bam files were analyzed together for structural variants using Delly v0.9.1 and Manta v1.6(Rausch et al. 2012; Chen et al. 2016). Both pipelines used a somatic filtering to filter out germline variants present in the germline. Resulting vcf files were analyzed using R and the library tidyverse.

### Optical Genome Mapping (OGM)

Optical genome mapping (OGM) requires at least 5mg of frozen tissue in order to extract at least 750ng of genomic DNA (gDNA). High-molecular weight DNA was extracted by the Bionano Prep SP Tissue Kit following the manufacturer’ instructions. DNA labeling was performed according to the DLS protocol with the DLE1 enzyme (CTTAAG sequence). The Saphyr Chip linearizes the labelled molecules and guides them into nanochannels to be imaged. Three samples were analyzed simultaneously for 3 days in order to obtain the highest possible coverage (Supp Table). Rare Variant Analysis (RVA) was performed using the Access software version 1.7 and solve version 3.7. The limits of detection of RVA are SVs > 5kb down to 5% allele fraction). A copy number variation tool running in parallel is able to detect CNVs > 500kb down to 10% allele fraction. Smap files generated by Bionano’s pipelines were downloaded and data analyses were performed with R using the tidyverse library.

The smallest size of the local captured SV is around 5kb. Resolution of breakpoints is ∼10 kb. Eligible tumor content in OGM was detected by (1) visible CNA or reported gains and/or losses in the cancer genome; (2) if no CNA reported the case need to be discarded if the number of SVs with VAF<0.4 is less than 20.

### Data and statistical analyses

To compare breakpoints of structural variants between WGS (A) and OGM (B) we used the distance Dist = |A_start – B_start| + |A_end– B_end|, where A_start is the starting chromosomal position of the variant detected by method A and A_end the ending chromosomal position of the same variant, and the same annotation is used for method B. The maximal distance between SVs junctions to call equivalent event was 50kb.

In SV analysis of WGS by Manta we filtered out all the variants supported by less than 10 reads, as well as deletions, duplications, insertions and inversions smaller than 1kb. Structural variants of the same type with distance between their junctions of less than 6kb were combined as a same event.

The Wilcoxon Mann-Whitney nonparametric test was used in order to assess the differences between the two groups of samples. P values < 0.05 were considered significant. The significance of the p values is annotated on graphs as * if p < 0.05, ** if p < 0.005 and *** if p < 0.001.

## Results

### Training series and quality control

Training set consisted in 27 TNBC (including 4 *BRCA1*-mutated, 1 *BRCA2*-mutated and 1 *RAD51C*-mutated cases), and 10 HGOC (including 1 *BRCA1*-mutated case) (Supplementary Table 1). 15 TNBC cases correspond to patients from the RadioPARP clinical trial (ClinicalTrials.gov Identifier: NCT03109080) (Loap et al. 2022). All cases from the training set were annotated as HRD or nonHRD (HRP Homologous Recombination Proficient) by shallowHRDv2 (Callens et al. 2023), based on shallow WGS (35 cases) or WGS (2 cases). All 37 cases had sufficient tumor content to provide unambiguous diagnostic by shallowHRDv2: 19 HRD (including the 2 cases with WGS) and 18 nonHRD. In addition, 8 HGOC were also tested by the Myriad MyChoice test (4 HRD and 4 nonHRD cases, consistent with the shallowHRDv2 status). Of note, OGM and WGS were performed from frozen blocks and shallowHRDv2 FFPE slides corresponding to different parts of the same tumor, explaining some differences in tumor contents and alterations between techniques.

Training series was tested by OGM with the coverage varying from 100X to 1400X. Common structural variants were filtered out using the OGM structural variants (SVs) database (RVA pipeline), as the corresponding germline sample was not analyzed. OGM characterized the training series by 10 types of SVs (see visual presentation of the typical cases in Fig 1 and more examples in Supp Fig 1). The total number of SVs reported were of wide range for 11 to 617 per sample.

**Figure 1.**
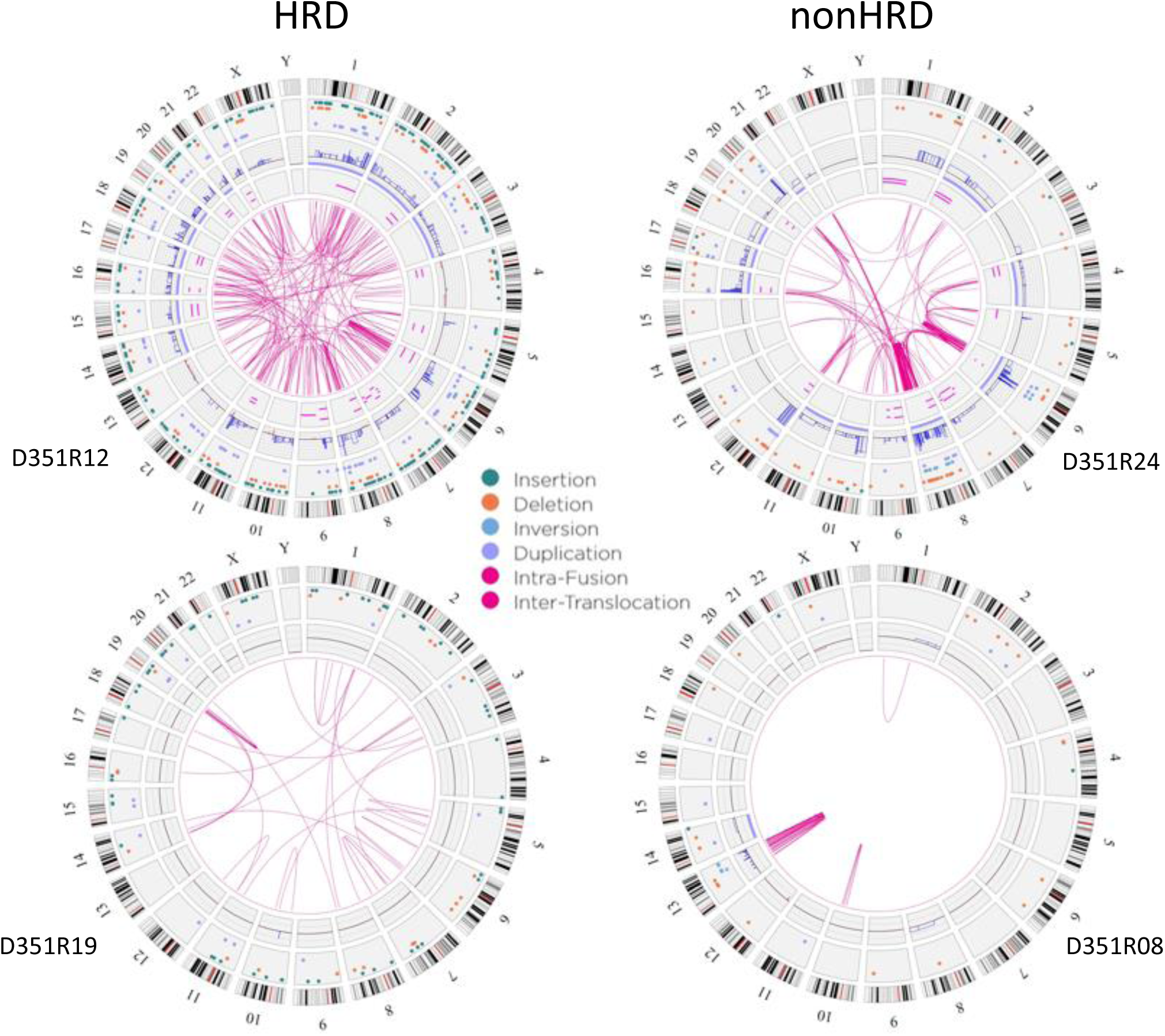
Circos plots of Optical Genome Mapping (OGM) of four representative examples of triple-negative breast carcinomas (TNBC) analyzed by RVA. Left panel: two examples of Homologous Recombination Deficient TNBC. Right panel: two examples of nonHRD (or HRP, Homologous Recombination Proficient) TNBC.

SVs are characterized by their variant allele frequencies (VAF) estimated by the RVA pipeline. The number of SVs with low VAF will depend on the coverage, as OGM uses a 5 molecules cut-off to call a SV. To diminish effect of marginal VAF in the samples with high coverage of over 500X, we filtered out SVs with VAF<0.01 in these samples.

### Tumor content

Five cases were suspected to have low tumor content from the visual estimation of the tumor sample by an expert pathologist and/or by the absence of copy number alterations (CNA) in the OGM report. Two of them were discarded as they carried the lowest numbers (11 and 33) of SVs detected by OGM, contrasting with numerous alterations displayed on the corresponding shallow WGS CNA profiles. The other three cases displayed >100 SVs, including one with the majority of SVs with VAF<0.1 (true low tumor content case). Thus, 35 cases were eligible for further analysis (18 HRD and 17 nonHRD).

### VAF dynamic in OGM output and subclonal events

High genomic coverage available with OGM approach allows getting correct results in low tumor content samples. From the other hand, high coverage in high tumor content sample can capture subclonal SVs, which can lead to over-estimation of the alteration load.

Typical violin plots of VAF in OGM output for TNBC or HGOC samples had wide peaks around 0.3 and significant amount of VAF>0.5, which might correspond to missed germline variants (not filtered by the OGM common SVs database) and/or true tumor SVs (Supp Fig 2A). We reasoned that SVs with 0.1<VAF<0.4 corresponded most probably to true clonal tumor SVs. High number of SVs with VAF<0.1 may evidence either low tumor content or subclones (Supp Fig 2A). We found 5 cases where the dynamic of VAF evidenced subclones: peak at 0.2<VAF<0.4, drop at 0.1<VAF<0.2 and extreme high peak at VAF<0.1 (Supp Fig 2A). Comparison with shallow WGS profiles confirmed that no detectable CNAs were found for SVs with VAF<0.1 (SVs of more 1Mb were considered to fit CNA resolution in shallow WGS). However, high number of SVs with VAF<0.1 might evidence relatively low tumor content, not subclones. To formally detect samples affected by subclonal admixture, we mapped the average VAF for SVs with 0.05<VAF<0.4 and the number of SVs with VAF<0.05 (Supp Fig 2B). Majority of cases follow the trend: the higher average VAF the lower number of SVs with low VAF. However, ∼15% of cases were out of this trend, representing cases with possible abundant subclonal SVs. These cases were flagged, as subclonal SVs can interfere with the upcoming analyses.

### Comparison of OGM and WGS in two BRCA mutated tumors

We analyzed two TNBC samples (one BRCA1mut and one BRCA2mut) by both OGM and WGS. Call for SVs in WGS was performed using Manta. Overall good correspondence was found for the BRCA2mut sample (∼70% of OGM and 45% of WGS were found in WGS and OGM, respectively); whereas correspondence was poor for the BRCA1mut sample (∼20% of OGM and 33% of WGS were found in WGS and OGM, respectively), due to low tumor content and relatively low WGS coverage compared to OGM (see Supp Table 2, Supp Fig 3 for details). This comparison suggested a higher sensitivity of OGM to capture SVs in low tumor content samples. We used the comparison of OGM with WGS to support re-annotation of SVs called by OGM.

### SV annotation

We redefined the OGM nomenclature of SVs, based on the known structural alterations characterizing TNBC and HGOC, and the actual comparison of the two cases with OGM and WGS, ending up with four main types of structural chromosomal aberrations (Table 1).

**Table 1.**
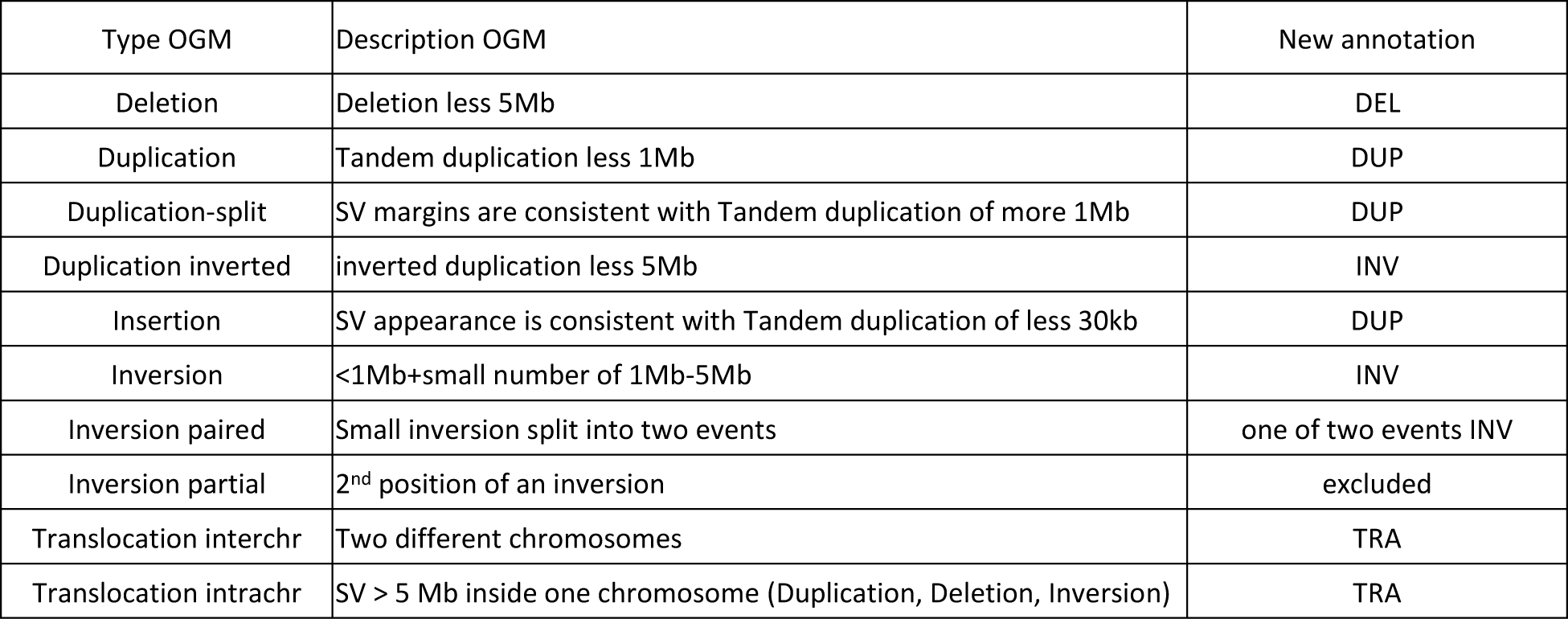
New annotation of Structural Variants detected by OGM pipeline.

OGM call three events associated with duplicated material: Duplication, Duplication-split and Duplication-inverted. Duplication corresponded to proven tandem duplication (TD) event and were subsequently denoted DUP. Duplication-split corresponded to a detected SV with boundaries compatible with a large TD of >1Mb size (Supp Fig 4A). Near 50% of TDs found in WGS corresponded to Duplication-split, so Duplication-split were classified as DUP (Supp Table 2). Duplication-inverted events were attributed to inversions (INV) because their quantity was rather correlated with that of inversions (R^2^=0.36) than DUP (R^2^=0) (Supp Fig 4B). INV would also aggregate Inversion-paired (as a single entity), while Inversion-partial were excluded. Insertions were classified as DUP, as most of Insertion events were consistent with TDs too small (<30kb) to be called as TDs by OGM, due to the resolution of the method, as previously reported (Mantere et al. 2021) and confirmed by WGS data (Supp Fig 4C, Supp Table 2). SVs designated by OGM as Inter-chromosomal and Intra-chromosomal translocations were classified as translocations (TRA).

### Selection of SVs significantly associated to HRD status

Given 35 OGM profiled cases characterized by 4 types of SVs (DEL, DUP, INV and TRA) and their respective HRD status, we aimed at feature selection for building a classifier that will allow testing HRD by OGM. Feature selection consisted in splitting DEL and DUP by size and mining TRA events to get the most significant association with HRD status (Table 2, Fig 2).

**Figure 2.**
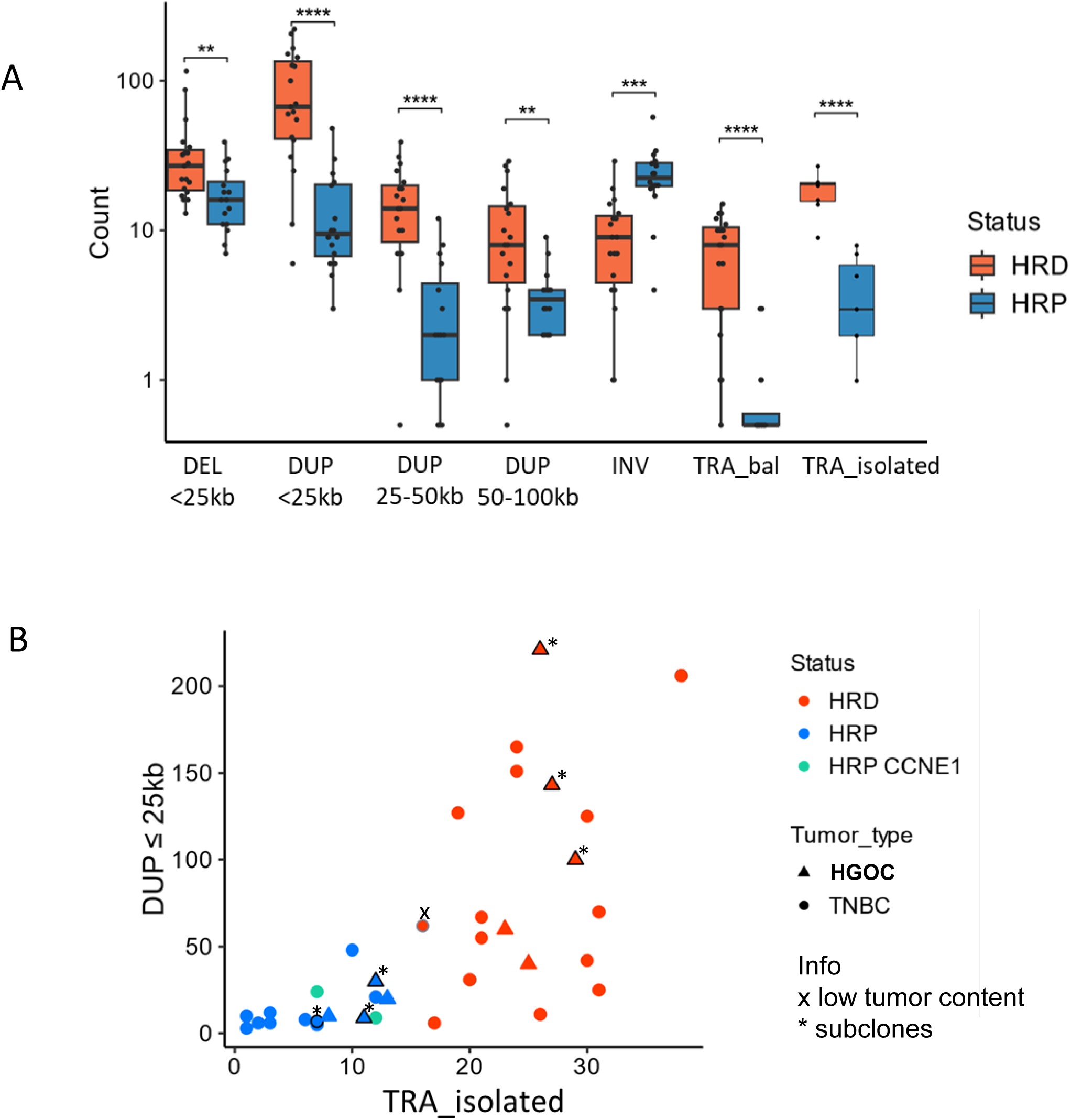
Differential analysis of structural aberrations called by OGM with respect to HRD status of tumors. A. Box plot representing the number of total structural variants with significant difference between HRD and nonHRD (HRP) tumors; translocation (TRA), isolated translocations (TRA_isolated), deletion (DEL), duplication (DUP) in HRD (orange) and HRP (blue) tumors; **, ***, **** indicate p_value <10^-2^, <10^-3^ and 10^-4^ respectively; Wilcoxon Mann-Whitney nonparametric test. B. 2D plot representing training series, with the number of isolated TRA (x-axis) and the number of Duplications less 25Kb (y-axis) as two the most differentiating HRD (red points) and nonHRD (blue points) features; additional information on low tumor content and subclonal admixture is indicated by “x” and “*”, respectively, at affected sample.

**Table 2.**
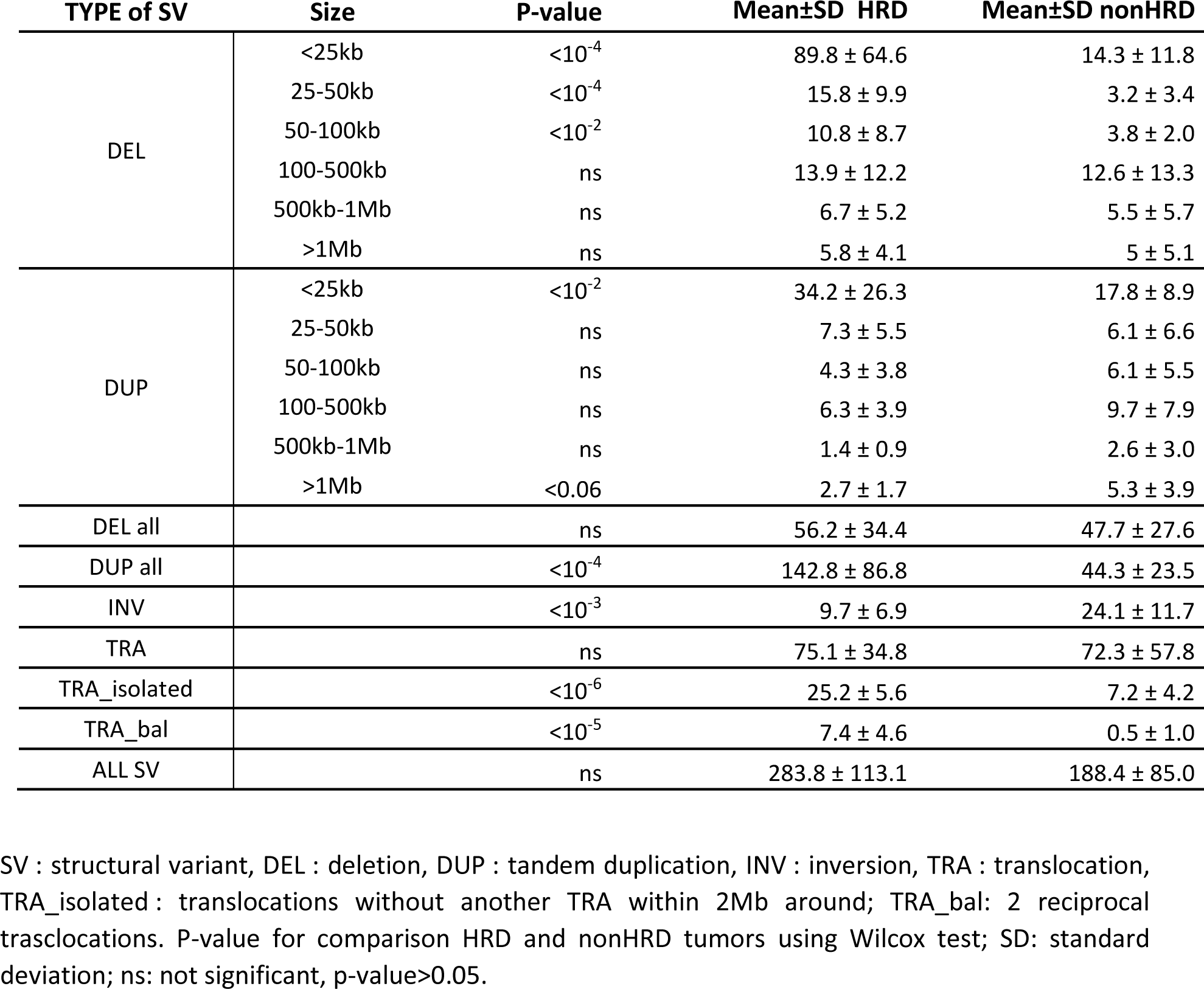
Statistics for structural variants (SV) and comparison of HRD (homologous recombination deficient) and nonHRD.

| TYPE of SV | Size | P-value | Mean±SD HRD | Mean±SD nonHRD |
| --- | --- | --- | --- | --- |
| DEL | <25kb | <10<sup>-4</sup> | 89.8 ± 64.6 | 14.3 ± 11.8 |
|  | 25-50kb | <10<sup>-4</sup> | 15.8 ± 9.9 | 3.2 ± 3.4 |
|  | 50-100kb | <10<sup>-2</sup> | 10.8 ± 8.7 | 3.8 ± 2.0 |
|  | 100-500kb | ns | 13.9 ± 12.2 | 12.6 ± 13.3 |
|  | 500kb-1Mb | ns | 6.7 ± 5.2 | 5.5 ± 5.7 |
|  | >1Mb | ns | 5.8 ± 4.1 | 5 ± 5.1 |
| DUP | <25kb | <10<sup>-2</sup> | 34.2 ± 26.3 | 17.8 ± 8.9 |
|  | 25-50kb | ns | 7.3 ± 5.5 | 6.1 ± 6.6 |
|  | 50-100kb | ns | 4.3 ± 3.8 | 6.1 ± 5.5 |
|  | 100-500kb | ns | 6.3 ± 3.9 | 9.7 ± 7.9 |
|  | 500kb-1Mb | ns | 1.4 ± 0.9 | 2.6 ± 3.0 |
|  | >1Mb | <0.06 | 2.7 ± 1.7 | 5.3 ± 3.9 |
| DEL all |  | ns | 56.2 ± 34.4 | 47.7 ± 27.6 |
| DUP all |  | <10<sup>-4</sup> | 142.8 ± 86.8 | 44.3 ± 23.5 |
| INV |  | <10<sup>-3</sup> | 9.7 ± 6.9 | 24.1 ± 11.7 |
| TRA |  | ns | 75.1 ± 34.8 | 72.3 ± 57.8 |
| TRA_isolated |  | <10<sup>-6</sup> | 25.2 ± 5.6 | 7.2 ± 4.2 |
| TRA_bal |  | <10<sup>-5</sup> | 7.4 ± 4.6 | 0.5 ± 1.0 |
| ALL SV |  | ns | 283.8 ± 113.1 | 188.4 ± 85.0 |

SV : structural variant, DEL : deletion, DUP : tandem duplication, INV : inversion, TRA : translocation, TRA\_isolated : translocations without another TRA within 2Mb around; TRA\_bal: 2 reciprocal translocations. P-value for comparison HRD and nonHRD tumors using Wilcoxon test; SD: standard deviation; ns: not significant, p-value>0.05.

The number and the size distribution in DUP captured by OGM with respect to HRD status showed clear difference (Supp Fig 5). Size distribution showed peak at ∼10kb and drops to almost zero at ∼25kb in HRD cases. Thus, we considered DUP_less25kb, DUP_25kb-100kb, DUP_100kb-500kb and DUP_more500kb as separate features. The most important feature associated to *BRCA1* inactivation is DUP_less25kb. By keeping DUP of other sizes, we anticipated TD phenotype or *CDK12*mut-associated TD phenotype, which were lacking in the training series.

Splitting DEL similarly to DUP, we noticed some enrichment of DEL_less 25kb in the two *BRCA2*mut cases (Supp Fig 5).

The number of TRA varied considerably in both HRD and nonHRD cases (Table 2). The highest number of TRA was found in few nonHRD cases, these TRA were clustered within few genomic regions with high density of SVs (Supp Fig 6A). We observed numerous TRA events sharing start, end or both, within the breakpoint resolution (denoted TRA_coupled). Part (but not all) of these coupled TRA events corresponded to reciprocal (or balanced; TRA_bal) translocations. TRA_single is characterized by a single junction without events closer 30kb. Among TRA_coupled, TRA_bal, TRA_single, only TRA_bal were significantly enriched in HRD.

Similar to the Large-scale genomic alteration (Popova et al. 2012), we introduced the notion of isolated TRA: TRA_isolated is TRA without another TRA closer than a given distance. This distance was fitted using the training series (Supp Fig 6B). A perfect discrimination between HRD and nonHRD cases was observed using distances from ∼1 to 2.5 Mb, demonstrating the robustness of this new feature. Using a 2Mb distance, we called TRA_isolated if no other TRA was found within 2Mb before or after, with TRA_bal counted as one TRA (Fig 2). As a cross-validation, we compared the number of TRA_isolated detected by OGM and of the large-scale genomic alterations (LGA) feature identified by the shallowHRDv2. These two features correlated well (R^2^ = 0.72; p_value<10^-4^, Supp Fig 6C), suggesting that the two independent methods are measuring a similar genomic feature.

TRA_isolated fully separated HRD and nonHRD in the training series with a gap: minimal number of TRA-isolated among HRD cases was found in low tumor content sample (TRA-isolated = 16) and maximal number of TRA-isolated among nonHRD cases were 14. We thus set up the threshold for HRD status to be TRA-isolated ≥ 16.

### Validation series

Validation series consisted of 12 TNBC (including 1 *BRCA1*-mutated, 1 *BRCA2*-mutated and 4 BRCAwt), and 18 HGOC (including 1 *BRCA1*-mutated, 2 *BRCA2*-mutated, 2 Myriad positive and 6 negative cases). ShallowHRDv2 was used for HRD diagnostics in 27 cases; 2 cases failed for tumor content, but were attributed to HRD (BRCA2mut) and to nonHRD (CCNE1 amplified, coincidence of HRD and CCNE1 amplification is possible but rare), respectively. One case with BRCA1 mut missing in shallowWGS was attributed to HRD.

OGM coverage range was similar to that of the training series (135-1460X). We performed the same workflow for validation series, including quality control and filtering SVs with marginal VAF. Four cases did not pass the quality control for tumor content. Altogether, the validation series included 26 cases with 10 HRD and 16 nonHRD status.

TRA_isolated were calculated and compared to the threshold defined from the training set. Five false HRD and no false nonHRD cases were found, including 2 cases with TRA_isolated close to the threshold and high numbers of TDs (including 1 case with *CCNE1* amplification), 2 cases over-passing the threshold by 3 and 4 (one with CCNE1 amplification) and one case with clear HRD phenotype. Thus, OGM revealed one case with true contradictory to shallowHRDv2 HRD diagnostics.

Overall, the large number of false positive cases in validation set is explained by higher diversity of nonHRD cases, which were not present in the training set. Using other selected features and more complicated recognition method can overcome this difficulty and is under development.

## Discussion

OGM analyses of breast or ovarian tumors allowed a cost effective and readily interpretable genome profiling, even in samples with low tumor content. In this study several informative SV features for HRD diagnostics were identified. The number of isolated inter- and intra-chromosomal translocation events, as defined by a free interval from another TRA event of around 2 Mb, represented a robust feature of HRD, by itself assuring high sensitivity. Incorporation of all informative features into a new classifier will help achieving higher specificity. Inclusion of *CCNE1*-amplification into tumor description and subsequent correction of the diagnosis could already reduce by two-fold false positive calls. OGM is thus an interesting and promising approach for HRD determination. More diverse validation series are needed for fine tuning HRD classifier based on OGM features. For future application of OGM in clinics, as pros, this method is cost-effective and highly efficient even with samples of low tumor content, it does not require complex analyses and high data storage. As cons, it requires a high quantity and quality of frozen tumor samples and an OGM dedicated platform. More technical developments and optimization are thus mandatory before realistic implementation of OGM in routine diagnosis of HRD.

## Acknowledgments

This work was supported by a Sponsored Research Agreement with Bionano Genomics and S.V. was supported by this grant. The authors thank François-Clément Bidard for his scientific and clinical inputs, the center of biological resources (CRB) of Institut Curie for providing frozen tumors fragments, in accordance with the ethical standards of the institutional and national research committees. DNA optical mapping was supported with the grant SESAME 2019 from the Region Ile France. The authors thank the NGS platform of the research center of Institut Curie. High-throughput sequencing was performed by the ICGex NGS platform of the Institut Curie supported by the grants ANR-10-EQPX-03 (Equipex) and ANR-10-INBS-09-08 (France Génomique Consortium) from the *Agence Nationale de la Recherche* (“*Investissements d’Avenir*” program), by the ITMO-Cancer Aviesan (*Plan Cancer* III), and by the SiRIC-Curie program (SiRIC grant INCa-DGOS-4654).

