## supplementary figure 3 for "Optical Genome Mapping for detecting Homologous Recombination Deficiency (HRD) in human breast and ovarian cancers"

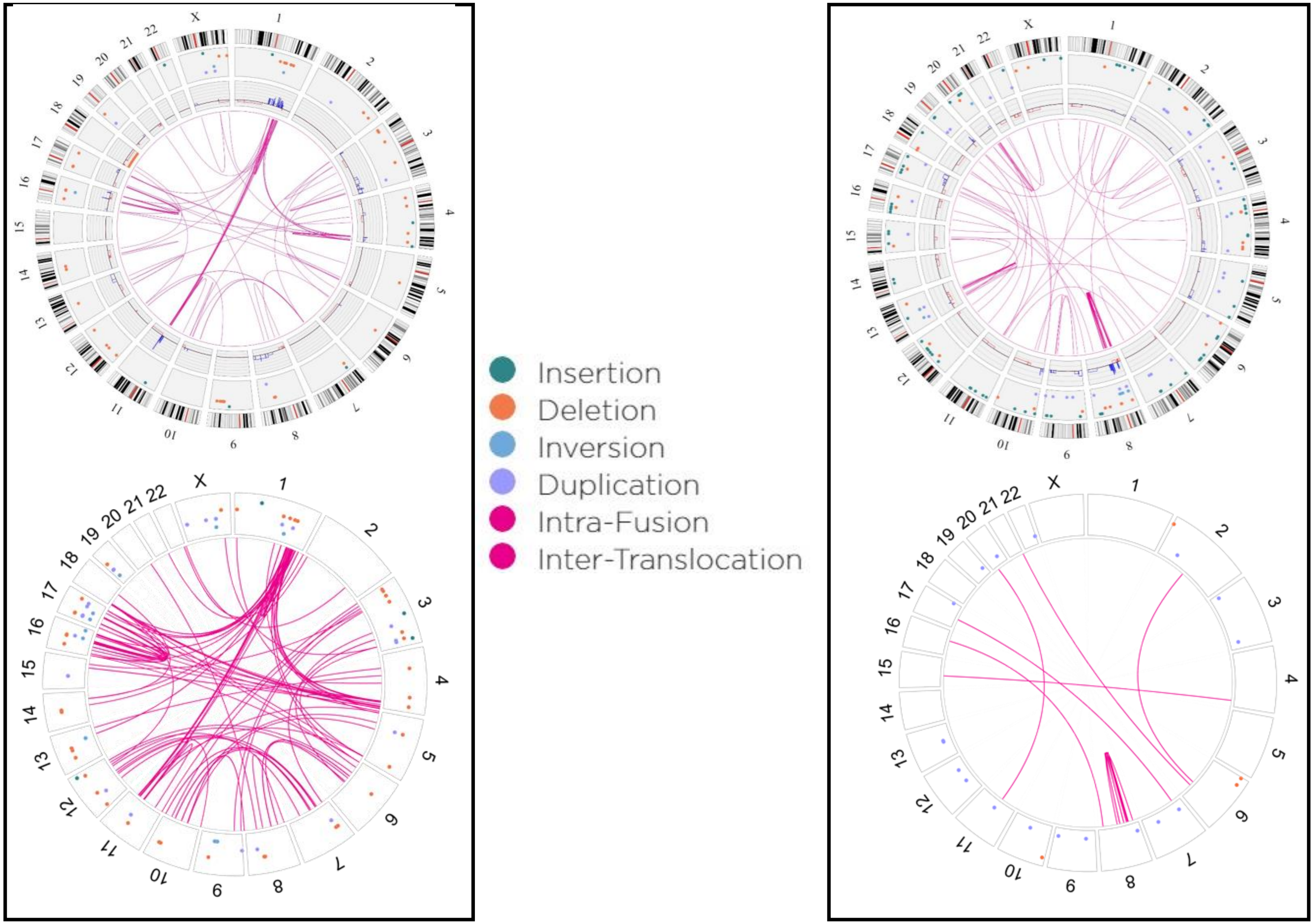


**Supplementary Figure 3.** Circos plots of OGM Structural Variant (SV) RVA analyses (top circos) and WGS SV analyses (bottom circos) in BRCA2-mutated (left panel) and BRCA1-mutated (right panel) triple-negative breast carcinomas (TNBC).
