## supplementary figure 4 for "Optical Genome Mapping for detecting Homologous Recombination Deficiency (HRD) in human breast and ovarian cancers"

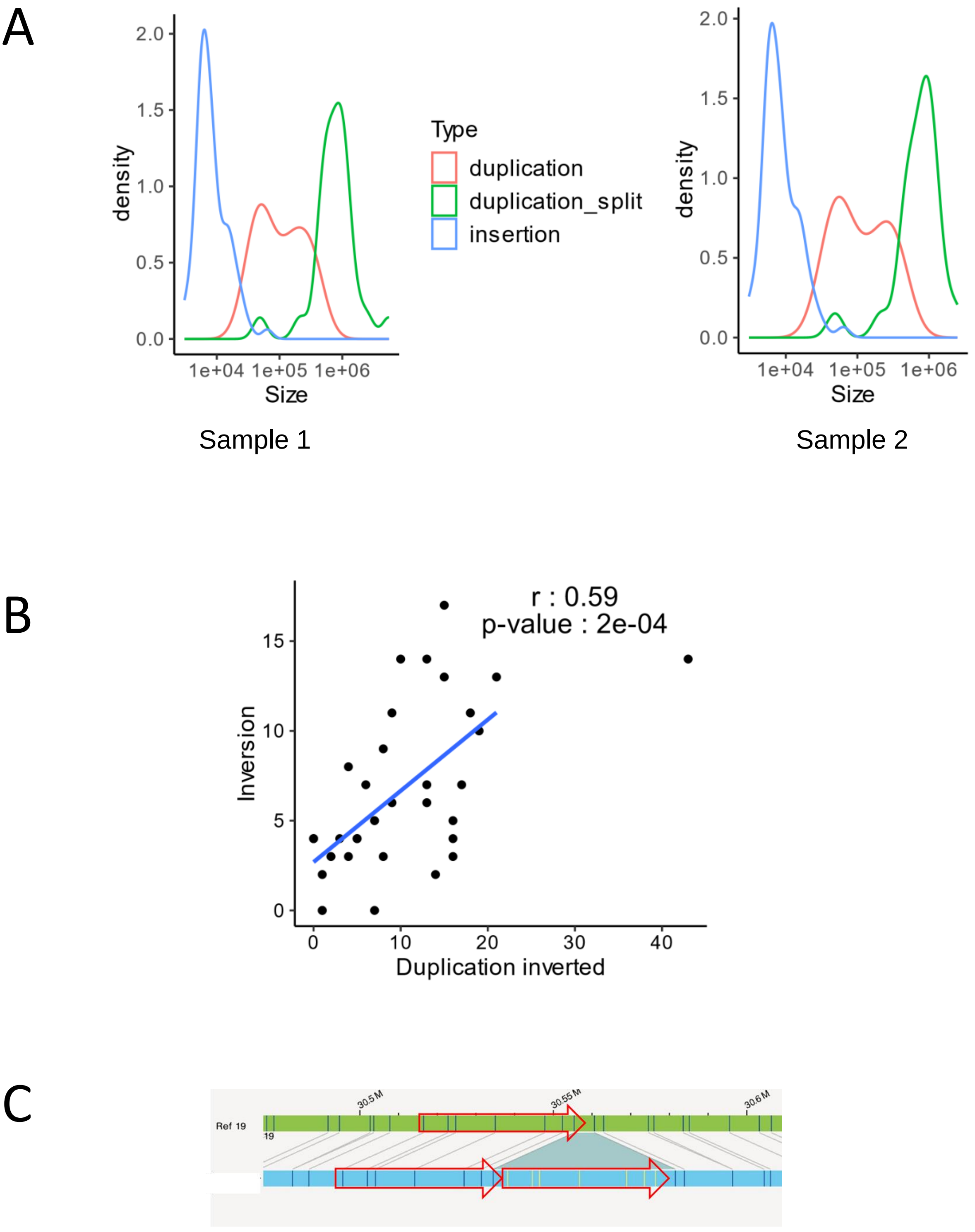


**Supplementary Figure 4. Annotation structural variants into main types: DUP and INV**
A. Density plot showing size distribution in Duplication, Duplication-split and Insertion events called by OGM pipeline. B. Correlation between the number of Inversions and Duplication-inverted called by OGM pipeline in the training series. C. Manual re-alignment showing that this rearrangement called INS by RVA could be re-classified as DUP.
