## Supplementary figures and images for "Optical Genome Mapping for detecting Homologous Recombination Deficiency (HRD) in human breast and ovarian cancers"

### supplementary figure 5

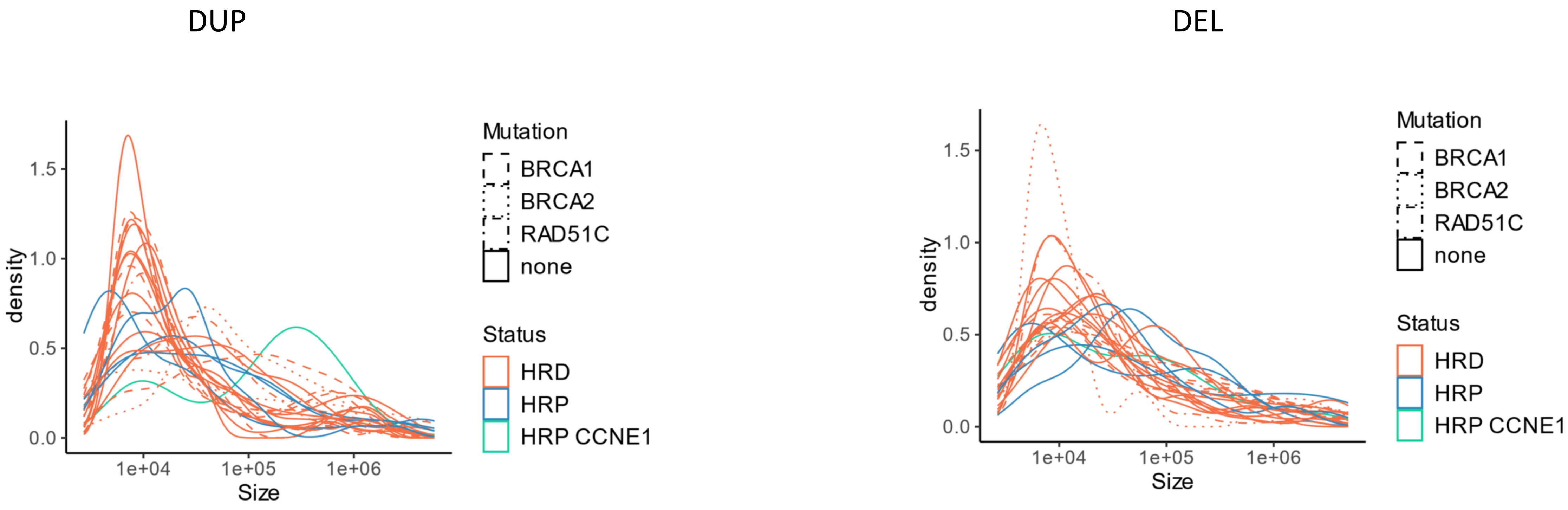


**Supplementary Figure 5.** Size distribution of DUP and DEL events in HRD and nonHRD (HRP) cases
