## supplementary figure 6 for "Optical Genome Mapping for detecting Homologous Recombination Deficiency (HRD) in human breast and ovarian cancers"

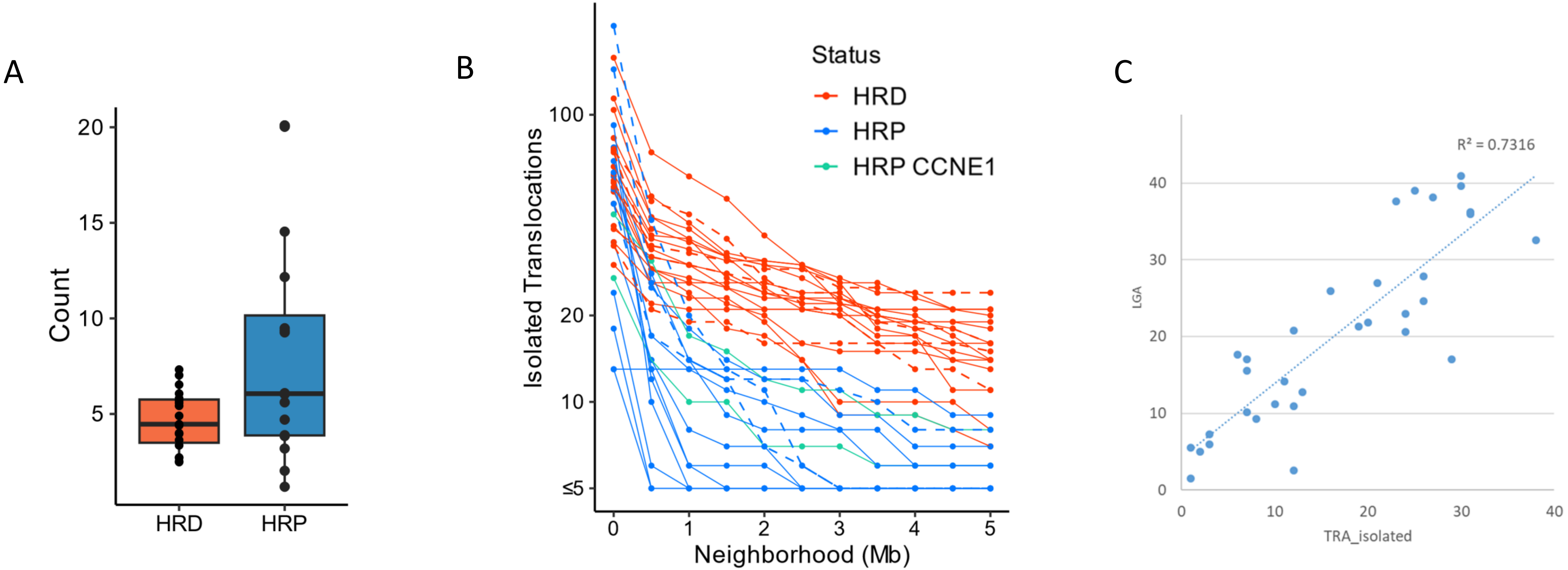


**Supplementary Figure 6. Mining translocations in HRD and nonHRD (HRP) tumors.** A. Number of TRA in 10% of the genome with the highest density of alterations. B. Number of isolated translocations depending on the distance to the TRA to be free of other TRA when calling TRA_isolated. C. Correlation between the number of isolated translocation found by OGM and the number of LGA (large genomic alterations) from shallowHRDv2.
