## supplementary table 1 for "Optical Genome Mapping for detecting Homologous Recombination Deficiency (HRD) in human breast and ovarian cancers"

**Supplementary Table 1. Training series for Optimal Genome Mapping (OGM)**

| Sample | Tissue | HR mutation | shallowHRDv2 | OGM coverage | Quality |
|---|---|---|---|---|---|
| D1090R22 | TNBC | RAD51C$^{GL}$ | HRD | 822 | |
| D1287E07 | TNBC | BRCA2$^{GL}$ | HRD | 454 | |
| D1287G07 | TNBC | none | nonHRD | 320 | discarded |
| D1309C01 | TNBC | none | HRD | 676 | |
| D351R03 | TNBC | none$^{GL}$ | nonHRD_CCNE1+++ | 509 | |
| D351R04 | TNBC | none$^{GL}$ | nonHRD | 488 | |
| D351R06 | TNBC | RAD51C$^{VUS}$ | HRD | 318 | |
| D351R08 | TNBC | none$^{GL}$ | nonHRD | 163 | |
| D351R09 | TNBC | BRCA1 | HRD | 532 | |
| D351R10 | TNBC | BRCA1$^{GL}$ | HRD | 411 | |
| D351R11 | TNBC | none$^{GL}$ | nonHRD | 407 | |
| D351R12 | TNBC | none | HRD | 788 | subclones |
| D351R13 | TNBC | BRCA1 | HRD | 290 | |
| D351R14 | TNBC | none | HRD | 813 | |
| D351R18 | TNBC | none$^{GL}$ | nonHRD | 740 | |
| D351R19 | TNBC | BRCA1$^{GL}$ | HRD | 276 | low tumor |
| D351R20 | TNBC | none$^{GL}$ | HRD | 359 | |
| D351R21 | TNBC | none | nonHRD | 800 | |
| D351R24 | TNBC | none | nonHRD | 568 | subclones |
| D844R35 | TNBC | | nonHRD | 1500 | |
| G1065B01 | TNBC | none | HRD | 241 | |
| G1079B01 | TNBC | none$^{GL}$ | nonHRD | 521 | |
| G1088A06 | TNBC | none$^{GL}$ | nonHRD | 741 | |
| G1088D06 | TNBC | | nonHRD_CCNE1+++ | 759 | |
| G962R06 | TNBC | | nonHRD | 429 | |
| WGS_01 | TNBC | BRCA1 | HRD | 940 | |
| WGS_02 | TNBC | BRCA2 | HRD | 718 | |
| D817R08 | HGOC | BRCA1 | HRD | 638 | |
| G834R01 | HGOC | none; M+ | HRD | 514 | |
| G834R04 | HGOC | none; M- | nonHRD | 1193 | subclobes |
| G834R09 | HGOC | M- | nonHRD | 305 | subclones |
| G847R05 | HGOC | M- | nonHRD_HER2+++ | 817 | |
| G852R08 | HGOC | none | nonHRD | 1101 | discarded |
| G873R02 | HGOC | none; M+ | HRD | 779 | |
| G873R07 | HGOC | none; M- | nonHRD | 1225 | |
| G876R05 | HGOC | none; M+ | HRD | 1458 | subclones |
| G925R10 | HGOC | none | HRD | 1292 | |

Tissue: TNBC triple negative breast cancer; HGOC high grade ovarian cancer;
HR mutation: mutation ($^{GL}$ germline, $^{VUS}$ variant of unknown significance) in Homologous Recombination (HR) pathway and/or Myriad myChoice status (M+ positive, M- negative)
shallowHRDv2: HRD (Homologous Recombination Deficiency) or nonHRD diagnostics obtained using shallow WGS using shallowHRDv2 pipeline
OGM coverage: mean whole genome coverage by genomic position (X)
Quality: results of quality control, including tumor content and subclones
