## supplementary table 2 for "Optical Genome Mapping for detecting Homologous Recombination Deficiency (HRD) in human breast and ovarian cancers"

**SAMPLE WGS_01**

| | | **OGM** | | | | |
|---|---|---|---|---|---|---|
| | | Deletion | Tandem duplication | BND between 2 chr | BND inside 1 chr | Total |
| **WGS** | Deletion | 26 (+5GL*) | | | | 44 |
| | Duplication | | 2 | | | 2 |
| | Duplication_inverted | | | | | 2 |
| | Duplication_split | | 2 | | | 3 |
| | Insertion | | 0 (+2 GL) | | | 9 |
| | Inversion | | | | 1 | 3 |
| | Translocation_interchr | | | 32 | | 45 |
| | Translocation_intrachr | 7 | 2 | | 9 | 23 |
| | Total | 44 | 9 | 103 | 24 | |

**SAMPLE WGS_02**

| | | **OGM** | | | | |
|---|---|---|---|---|---|---|
| | | Deletion | Tandem duplication | BND between 2 chr | BND inside 1 chr | Total |
| **WGS** | Deletion | 3 (+10 GL) | | | | 33 |
| | Duplication | | 1 | | | 11 |
| | Duplication_inverted | | | | | 4 |
| | Duplication_split | | 10 (+3 GL) | | | 29 |
| | Insertion | | 6 | | | 75 |
| | Inversion | | | | | 10 |
| | Translocation_interchr | | | | 7 | 33 |
| | Translocation_intrachr | 1 | 4 | | 6 | 35 |
| | Total | 6 | 23 | 66 | 15 | |

*GL: germline

**Supplementary results**

We compared SVs detected in WGS by Manta and in OGM by the RVA for two samples.

Overall good correspondence was found for one sample. For the other, correspondence was minor probably due to low tumor content in the sample and lower coverage of 70X in WGS compared to the 720X in OGM. Of note, SVs that are detected multiple times due to small label differences in the molecules were filtered out; some events in both samples were not detected by Manta but were visible by IGV.

For **translocations**, 33/45 and 7/33 of OGM's interchromosomal translocations are also detected by Manta. Of note, 7 others from the first sample correspond to cumulative translocations detected in WGS not identified by OGM due to the low resolution. The length of the unmapped in-between translocation part can be found in OGM's data, but the annotation of interchromosomal translocation remains the same. We will not consider those events differently.

Pairs of translocations detected by OGM at almost the same positions corresponding to the opposites part of the chromosomes (left or right of the breakpoint) are present in both samples (2 and 1). Those events correspond to balanced translocations. We will add a subcategory in the translocations to represent them.

Intrachromosomal translocations correspond to deletions, duplications and breakends all larger than 5Mb detected by Manta (respectively 7,2,9 out of 23 for sample1 ; 1,4,6 out of 36). Thus, the SV class intrachromosomal translocations represents large rearrangements inside one chromosome. ("Intrachromosomal fusion breakpoints involve regions typically (but not always) at least 5 Mbp away from each other on the same chromosome" - Bionano-Solve-Theory-of-Operation-Structural-Variant-Calling)

For **deletions**, 26/44 and 3/33 of OGM's deletions are also detected by Manta. Out of the remaining ones, there are respectively 5 and 10 that are germline variants not present in the control database of Bionano. *(VAF from 0.28 to 0.98 sample1 ; 0.26 to 0.68 sample2)*

For **other SVs**, OGM detected 3 different categories of duplications: the duplications, the duplication split and the inverted duplication. For the duplications, 2/2 and 1/9 are detected by Manta as tandem duplications (TD). For the duplication split 2/4 and 10/31 are detected by Manta as TD. Moreover, 1 of the remaining ones in both samples correspond to duplications already detected by OGM. They are thus duplicates that need to be removed. For the inverted duplications, none out of the 2 and 4 are detected.

OGM also detects insertions. Since this is not an SV type detected by Manta and that it has previously been proven that insertions smaller than 30kb are called duplications because the label density may not be informative enough to exactly determine the origin of the inserted material (ref Mantere et al 2021), we have compared them to TDs found by Manta. For both samples we found 0/9 and 6/76 OGM insertions corresponding to Manta's TD. In the remaining ones, 2 and 3 are germline TDs, and 1 in sample 2 is a duplicate of an OGM duplication already detected.

Theses comparisons suggest that duplications, insertions and duplications split are all tandem duplications. Moreover, looking at the size of those events in both samples, we can see that those three categories have different sizes, insertions being the smallest and duplication split the largest. Thus insertions are too small, and duplication split too large to be detected as duplications by OGM. However, we cannot exclude the possibility that some insertions are inverted duplications. Further, we will consider them together as one SV category, and we will remove the duplicates.

OGM detects inversions in different categories: inversion_paired that represent in pairs the different positions of one inversion, inversions and inversion_partial that represent the other positions of the inversions. We will only keep the inversions and remove their paired inversion_partial, and one of the paired inversions. Comparison of those inversions lead to 1/3 and 0/10 of them that are detected as BND. Since inversions and inverted duplications represent a similar event, we will group them together for further analysis.
