## supplementary figure 1 for "Optical Genome Mapping for detecting Homologous Recombination Deficiency (HRD) in human breast and ovarian cancers"

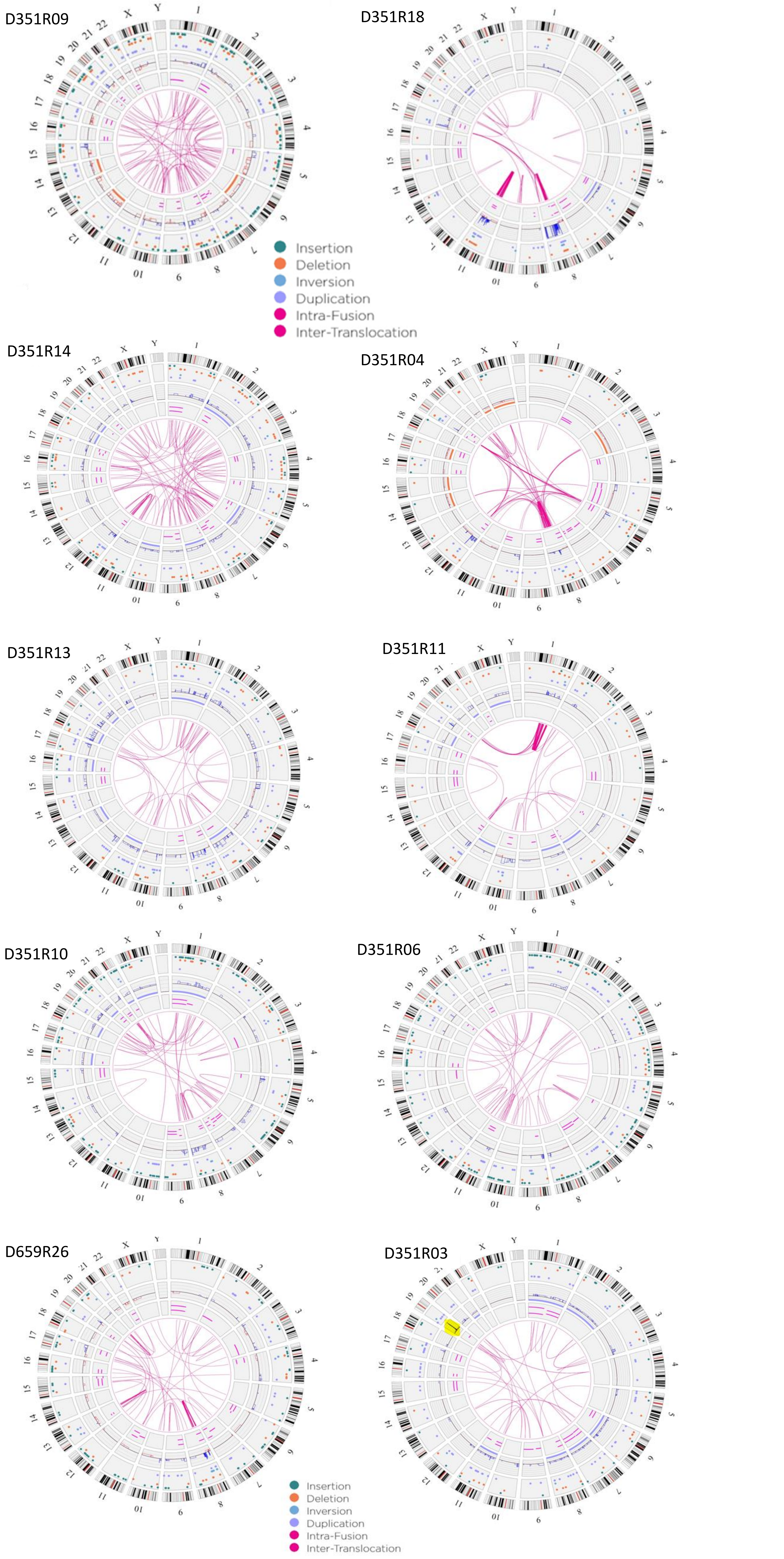


**Supplementary Figure 1.** Circos plots of Optical Genome Mapping (OGM) of 10 triple-negative breast carcinomas (TNBC) analyzed by RVA.
