## supplementary figure 2 for "Optical Genome Mapping for detecting Homologous Recombination Deficiency (HRD) in human breast and ovarian cancers"

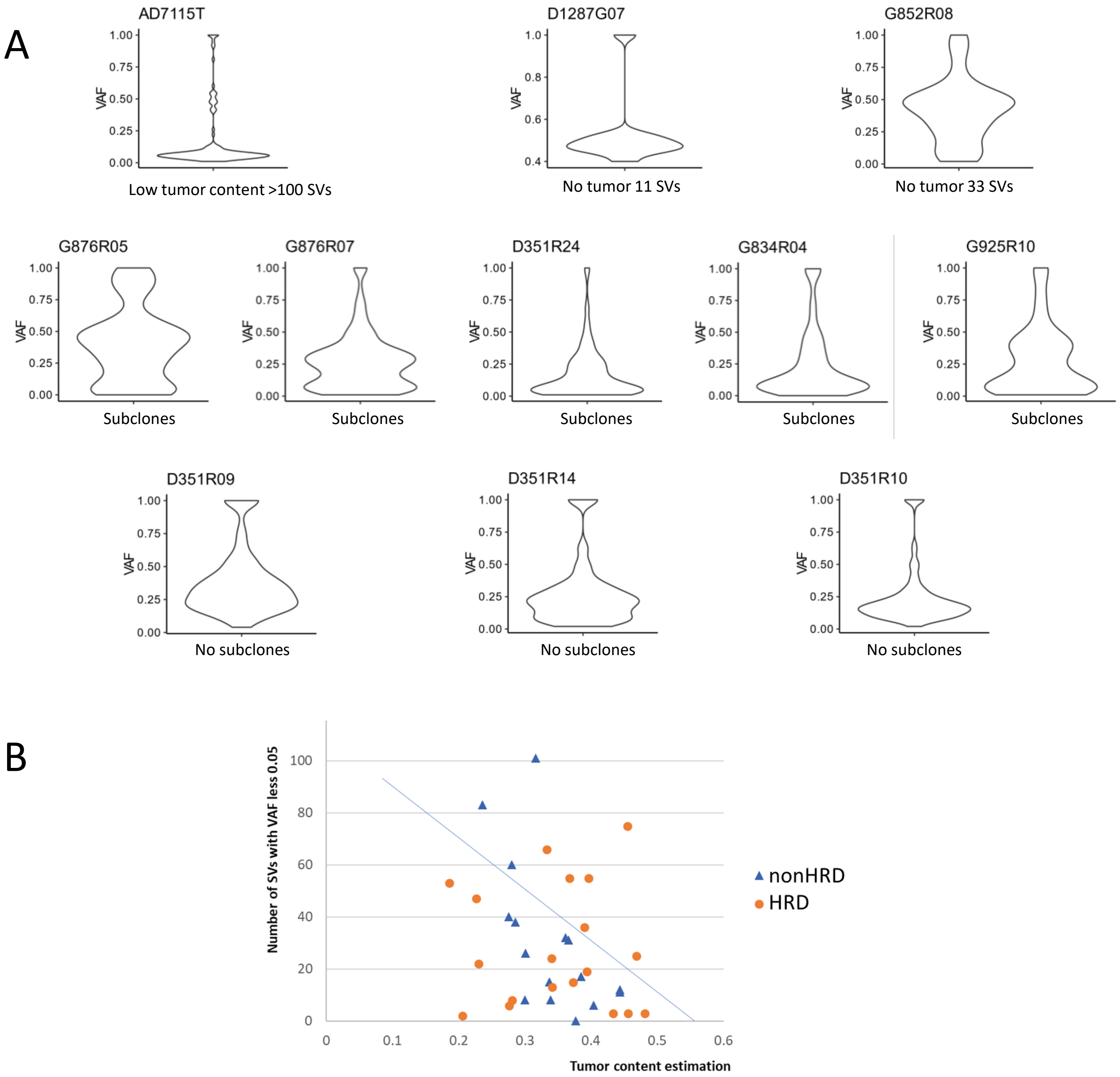


**Supplementary Figure 2.** VAF distribution, estimation of the tumor content and subclonal admixture in OGM delivery. A. Violin plots of VAF distribution for the selection of samples, top row: low tumor content; middle row: subclones; bottom row: no subclones. B. Samples are plotted by their mean VAF (0.05<VAF<0.4) multiplied by 2 (x axis) and the number of SVs with VAF<0.05 (y axis); outliers (above the trend line) could be considered as the cases enriched with subclones.
